# Plantar flexor deficits following Achilles tendon rupture: a novel small animal dynamometer and detailed instructions

**DOI:** 10.1101/2022.08.01.502343

**Authors:** My M. Tang, Courtney A. Nuss, Natalie Fogarty, Josh R. Baxter

**Affiliations:** Department of Orthopaedic Surgery, University of Pennsylvania, Philadelphia, Pennsylvania, USA

**Keywords:** Dynamometry, preclinical model, orthopaedic surgery, rehabilitation, rat, function

## Abstract

Plantar flexor functional deficits measured using joint dynamometry are associated with poor outcomes in patients following Achilles tendon rupture. In this study, we developed a small animal dynamometer to quantify functional deficits in a rat Achilles tendon rupture model. Like our reported plantar flexor deficits in patients recovering from Achilles tendon ruptures, we found in our small animal model functional deficits across the ankle range of motion, resulting in an average 34% less positive work being done compared to the uninjured contralateral limb. These functional deficits are similar to 38% less plantar flexor work done by patients who were treated non-surgically in our prior research. Further, these deficits were greater in plantar flexion than dorsiflexion, which agree with clinical complaints of limited function during tasks like jumping and hiking. These findings highlight the impact of muscle-tendon loading during early tendon healing on long-term plantar flexor function and serve as compelling evidence that our Sprague Dawley rat model of an Achilles tendon rupture recapitulates the human disease. We provide thorough documentation for other groups to build their own dynamometers, which can be modified to meet unique experimental criteria.

**SIGNIFIGANCE:** Preclinical models are critical tools for translating knowledge discovery to clinical decision making. We developed a low-cost and flexible joint dynamometer that quantifies joint function in small animals. Here, we used a rat model to test the implications of Achilles tendon ruptures not surgically repaired on plantar flexor function. We found that Achilles tendon ruptures in a rodent model closely resemble the functional deficits our group has observed in patients.

## Introduction

Quantifying functional deficits following musculoskeletal injury are a cornerstone of clinically impactful research. Joint dynamometry has long been considered the ‘gold-standard’ for quantifying joint-level function in patient populations (Dvir and Müller, 2020; Martin et al., 2006; Ushiyama et al., 2017). Despite these established techniques in patient populations, there is less consensus on how to quantify functional deficits in pre-clinical models that aim to recapitulate human disease conditions. In this short communication, we describe a flexible, low-cost, and convenient small animal dynamometer that quantifies joint-level functional capacity.

We then use an established rat Achilles tendon rupture model (Freedman et al., 2016) to demonstrate functional deficits that show strong similarities to our observations in patient populations (Baxter et al., 2018; Hullfish et al., 2019).

## Methods

We designed this small animal dynamometer to be flexible to accommodate a range of animal species and joints tested. The dynamometer has 8 primary components (**Figure 1**): 1) a platform constructed from 2 aluminum breadboards with a grid of threaded holes to secure instrumentation and 3D printed parts, 2) an instrumented foot plate assembly comprised of a torque cell in series with a servomotor and 3D printed foot plate, 3) 3D printed parts to safely secure the test animal to the dynamometer, 4) a torque cell signal conditioner to transform the torque cell measurements into an analog signal, 5) a data-acquisition board to measure the torque cell converted signal and send a command signal to the isolated constant current stimulator, 6) a microcontroller to send position commands to the servomotor, 7) an isolated constant current stimulator to elicit controlled muscle contractions, and 8) a graphical user interface to run the testing protocol and record the experimental data. Here we briefly describe each component and provide greater detail – including mechanical drawings, parts info, software files, and dissection protocols – in the supplemental materials and publicly posted repositories.

**Figure 1.**
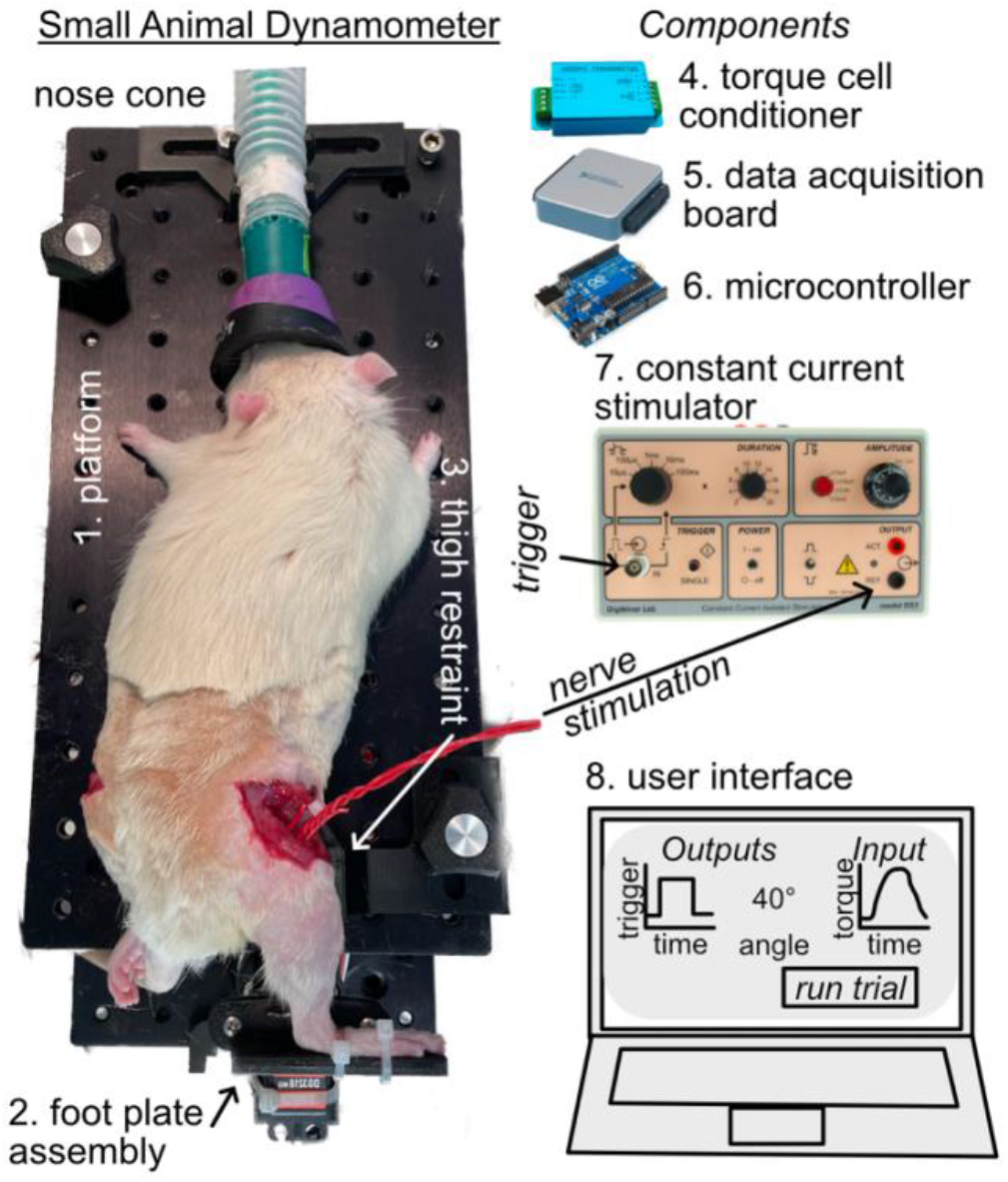
We constructed our small animal dynamometer (*left*) using a (1) pair of aluminum breadboards, a (2) foot plate assembly, and a (3) thigh restraint to position the ankle and measure reaction joint torques. We used electrical components (*right*) to send and receive experimental data. We used a (4) torque cell conditioner to transform the ankle reaction torque that was read by a (5) data acquisition board and a (6) microcontroller to control the position of the ankle. We elicited maximal plantar flexor contractions using a (7) constant current stimulator that was controlled by a (8) custom-developed user interface.

Our small animal dynamometer prescribes static joint angles and measures maximal isometric muscle contractions elicited by sciatic nerve stimulations. By measuring these maximal joint moments across a range of joint postures, we reliably quantify the joint-moment relationship. This relationship is the joint-level equivalent of the force-length properties of skeletal muscle (Maganaris, 2001). We decided to quantify joint function via isometric dynamometry as opposed to isokinetic dynamometry for several reasons: 1) our published data suggest that patient function is limited in plantar flexion when the shorter muscle fascicles in series with an elongated tendon are incapable of doing positive work (Baxter et al., 2018), 2) other functional tasks like walking result from isometric plantar flexor contraction (Fukunaga et al., 2001) rather than high velocity contractions, and 3) based on these physiologic considerations, we selected a low-cost servomotor that does not have the performance characteristics to prescribe joint rotations at sufficiently high speeds. However, modifying our small animal dynamometer to accommodate a high-speed servomotor is possible.

We performed a series of experiments to demonstrate the clinical utility of our small animal dynamometer. In this set of experiments, 9 adult male Sprague Dawley rats (weight: 428+\- 40 g) underwent a surgically induced Achilles tendon rupture, followed by 2-weeks of joint immobilization in plantar flexion, and then 2-weeks of unrestricted cage activity. These procedures were approved by our Institutional Animal Care and Use Committee and described elsewhere in greater detail (Freedman et al., 2016). Briefly, we made a small incision near the Achilles tendon while the animal was anesthetized using 2% isoflurane. Then, we resected the plantaris longus tendon and severed the mid-portion of the Achilles tendon using the blunt edge of an 11-blade. After closing the skin with an absorbable suture (Vicryl 9-0, Ethicon), we immobilized the ankle in plantar flexion using a straight plastic splint that put the ankle near 90 degrees of unloaded plantar flexion. We used cotton and self-adhesive wraps to hold the leg in that position and poured bone cement (polymethyl methacrylate) to prevent the animals from chewing off the brace. We checked animals daily and changed casts as needed. After 2-weeks of immobilization in plantar flexion, we removed the braces and provided the rats with unrestricted cage activity to simulate return to full weight bearing in patients.

We implanted a pair of 24-gauge electrical wires around the sciatic nerve to maximally stimulate the plantar flexor muscles (**Supplemental Figure S1**). To do this, we anesthetized each animal using isoflurane 1-4% and maintained body temperature using a circulating hot water pad during the sciatic nerve dissection and a heat lamp during the dynamometer testing. We shaved the hindlimbs of each animal and made a small skin incision on the lateral aspect of the thigh parallel with and just caudal to the femur using scissors. To gain exposure to the sciatic nerve, we used blunt dissection to separate the vastus lateralis and biceps femoris muscles. We carefully dissected the connective tissue surrounding the nerve and placed electrode wires around the nerve. We then filled the surgical window with warmed paraffin oil, which acts as an insulator while preventing the nerve from drying out. We repeated this procedure on both legs so that we could analyze contralateral ankle function.

We tested a wide range of ankle postures with the knee held in 30 degrees of flexion to quantify plantar flexor function. During pilot testing, we determined that 20-degree increments centered around a neutral ankle position extending to 60 degrees dorsiflexion (30 degrees) and 60 degrees plantar flexion (150 degrees) fully characterizes the physiologic range of ankle motion in the rat (Freedman et al., 2016). We tested these 7 ankle positions with 15 seconds of rest over 3 sets of contractions. During pilot testing, we found that 15 seconds of rest was sufficient to minimize fatigue (**Supplemental Figure S2**) while facilitating time efficient experiments. We randomized the order of each ankle position within a given contraction set to minimize the effects of fatigue. We then delivered 500 ms square wave stimulations (100 hz, 100 µs pulse width, 1-3 mA) using a constant current stimulator (DS3, Digitimer, UK). Before each testing session, we performed isometric contractions with the foot neutrally aligned to set the stimulation amplitude. Because our research is focused on plantar flexor function, we transected the dorsiflexor tendons prior to eliminate the effects of co-contraction. We randomized the ankle order to eliminate potential effects of testing sequence bias. We then calculated the peak plantar flexor moment generated at each ankle position and the plantar flexor work done across the range of motion. We hypothesized that the Achilles tendon rupture would reduce plantar flexor moment across the range of motion and lead to less plantar flexor work based on our earlier patient findings (Hullfish et al., 2019). Therefore, we tested for these differences using one-tailed t-tests with α = 0.05.

## Results

The Achilles tendon rupture caused plantar flexor functional deficits across the ankle range of motion (**Figure 2**). When normalized to the contralateral uninjured ankle, we found that these functional deficits were most pronounced in plantar flexion. Peak plantar flexor moments were 29-34% less between 60 degrees dorsiflexion to 20 degrees plantar flexion (p<0.002). These deficits increased to 41% in 40 degrees plantar flexion and to 48% in 60 degrees plantar flexion. These functional deficits across the ankle range of motion explain a 34% decrease in plantar flexor work (p<0.001) in the injured limb (301.4 ± 65.5 N mm) compared to the control limb (465.9 ± 62.7 N mm).

**Figure 2.**
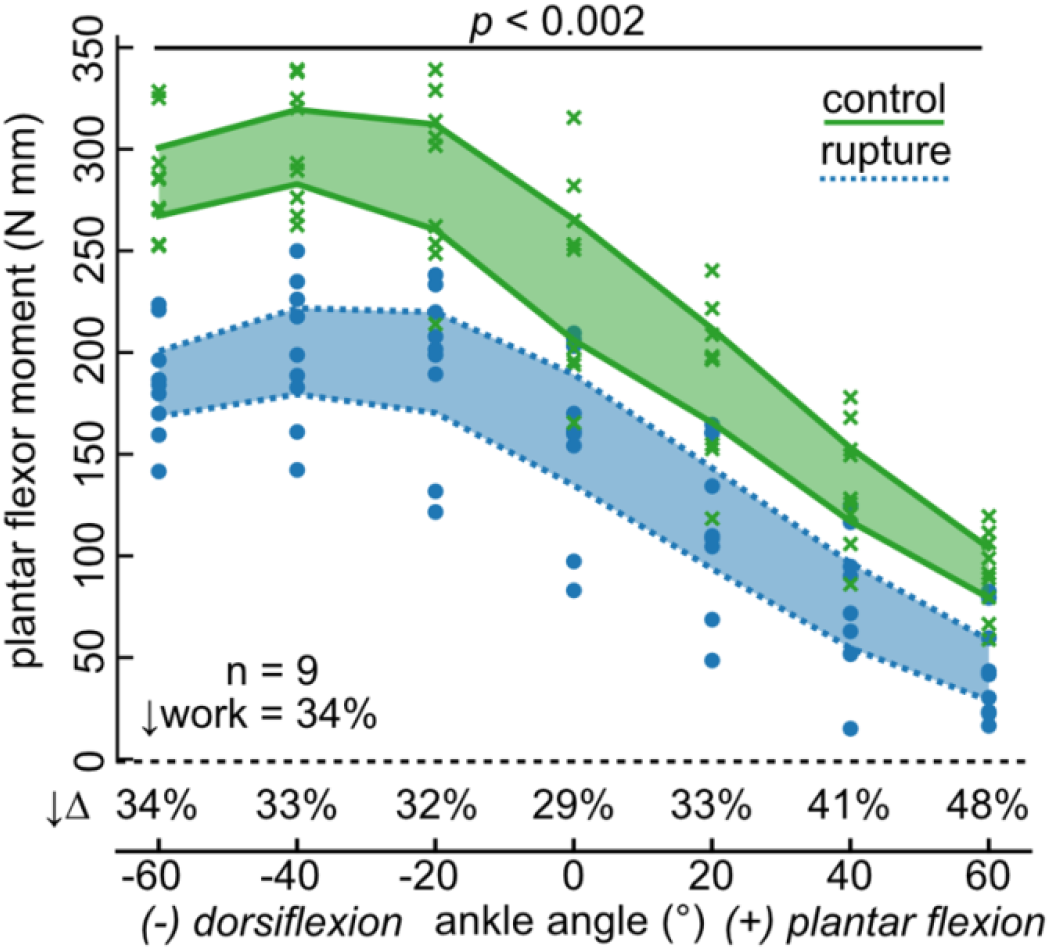
Plantar flexor moments were on average 35% less following Achilles tendon rupture compared to the uninjured contralateral limb. When normalized to the contralateral ankle moments, these functional deficits were most severe in 40 and 60 degrees of plantar flexion. Shaded areas are 95% confidence intervals.

## Discussion

We developed this small animal dynamometer to quantify plantar flexor functional capacity in a rat Achilles tendon injury model. We selected geometries, motor size, and torque cell range that met these experimental criteria. However, each of these components can easily be replaced or modified to meet the demands of other small animals or joints. For example, testing a larger animal like a rabbit could be done by using a torque cell with greater measurement range and a servomotor capable of generating greater torque. While developing this system, we found the most challenging aspect was establishing a method to reliably secure the animal to the dynamometer. After several iterations of testing, we decided to secure the thigh to a 3D printed knee bracket that positions the knee at 30 degrees of flexion using a plastic cable tie. Because our dynamometer testing is currently an end-of-life procedure, we used an invasive technique that bluntly dissection between the femur and knee flexor muscles to route the cable tie. However, during pilot testing we found non-damaging solutions were also effective – albeit less secure – in holding the knee at the prescribed 30 degrees of flexion.

Preclinical animal models can be powerful research platforms to test a wide range of questions in the pursuit of improving patient care. However, validating these models with patient conditions should consider clinically relevant – and preferably, clinically implemented – functional outcome measures. Using our small animal dynamometer, we confirmed that a preclinical rat model (Freedman et al., 2016; Hillin et al., 2019; Huegel et al., 2019; Leahy et al., 2022) is a valid and relevant model to study the effects of Achilles tendon ruptures on plantar flexor function that directly translate similar measurements made in patients (Baxter et al., 2019, 2018; Hullfish et al., 2019; Silbernagel et al., 2012). The angle-torque curves we measured in our non-surgically repaired rats compare favorably to the same measurements of plantar flexor function made in a cohort of patients treated non-surgically for an acute Achilles rupture (**Figure 3**). Non-surgically repaired Achilles tendon ruptures in rats caused a 34% deficit in positive plantar flexor work 4-weeks after the injury while the same injury caused a 38% deficit in our patients 14-weeks after the injury (Hullfish et al., 2019). Integrating functional joint assessments into preclinical research using a small animal dynamometer may prove to be a critical step in translating findings to clinical practice. An important translational aspect of our joint dynamometer is the similarities in both the shape and scale of plantar flexor deficits between preclinical and clinical populations. Because many physical therapy clinics utilize joint dynamometry for routine patient care and screening, we believe this paradigm can bridge preclinical and clinical research to maximize the translational impact of preclinical research on patient care.

**Figure 3.**
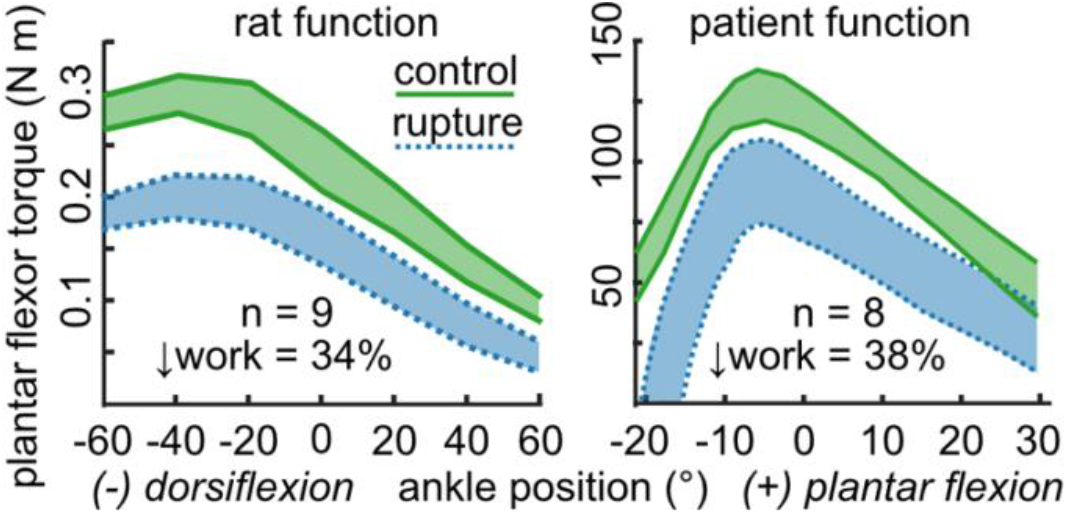
Plantar flexor deficits were similar in our small animal experiments and patient observational study (Hullfish et al., 2019) where both cohorts experienced Achilles tendon ruptures and received non-surgical treatment. Despite the much larger range of ankle motion in the rat, both species experienced similar deficits in plantar flexor work done measured using dynamometry.

Our dynamometer paradigm has several limitations that should be carefully considered when deciding how to quantify joint function in a small animal experiment. The system invokes involuntary maximal isometric contractions by stimulating the sciatic nerve. While this fully characterizes joint-level functional capacity, it neglects the potential effects of pain and neural inhibition on plantar flexor function. After correcting for differences in body weight, our study found peak plantar flexor torques measured at a neutral ankle angle were approximately 10% less than earlier reports (Warren et al., 2004). We suspect that differences in animal weight and joint posture explain the variations between these two studies. We decided to use isometric contractions across a range of joint postures to quantify joint torque-angle properties instead of continuous isokinetic contractions. There are certainly experimental questions that benefit from quantifying force-velocity capabilities and future development should focus on implementing a higher performance servomotor. Additionally, stimulating the sciatic nerve excites the smaller plantar flexor muscles in addition to the larger muscles of the posterior compartment. Based on our clinical question, we determined that quantifying the complete plantar flexor capacity was most reflective of our assessments in our patient population (Baxter et al., 2019, 2018; Hullfish et al., 2019; Silbernagel et al., 2012). Further, we found that dissecting the sciatic nerve was simpler and lower risk than dissecting the smaller tibial nerve. However, our dynamometer paradigm is agnostic to which nerve is stimulated and we expect it is robust for assessing any type of isometric joint contraction.

In conclusion, we developed and used a clinically relevant small animal dynamometer to assess plantar flexor functional deficits following an Achilles tendon rupture. Using this approach, we provide compelling evidence that our Sprague Dawley rat model of an Achilles tendon rupture closely recapitulates the human disease. In hopes of supporting other research groups, we provide supplemental and publicly posted files for others to build and use our dynamometer for their own research.

## ACKNOWLEDGEMENTS

We thank Mr. Todd Hullfish, Mr. Hammo Ahmad, and Ms. Liala Sofi for assisting in the small animal surgeries and immobilization protocol.

