## Supplemental Material for "Plantar flexor deficits following Achilles tendon rupture: a novel small animal dynamometer and detailed instructions"

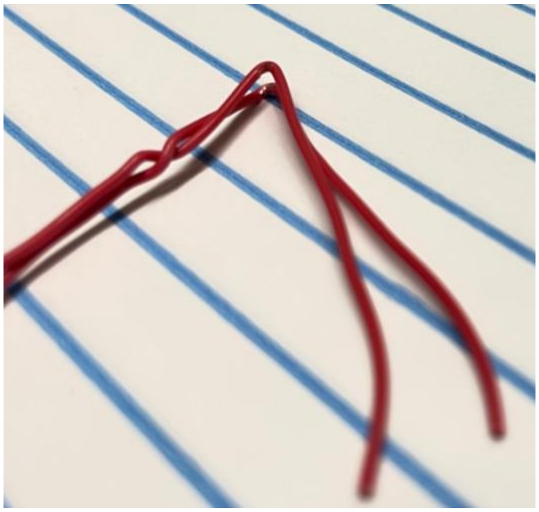

**Figure S1.** We carefully shaved the insulating coating of 24 AWG wires to expose the bare electrode using a scalpel. By folding the electrode pair at this cut location, the electrodes can be placed around the sciatic nerve.

**Nerve electrode.** We tested several iterations of nerve electrodes before finally finding a design that we considered to be the most robust, reliable, and easy to fabricate and use. This final design used 24 AWG single strand insulated wire. We cut 2 lengths of ~16” and twisted them together at several spots along the length of the wire to prevent the wires from moving with respect to one another. We then folded both wires approximately 1” from the end of the wire pair. We then cut off a small amount of insulating material to expose the bare wire (**Figure S1**). These small, exposed areas of the wire are what contact the sciatic nerve. After placing the nerve electrode, it is important to fill the space with warmed paraffin oil to keep the nerve and surrounding soft tissues hydrated. We want to stress several things for reliably using this nerve cuff setup: 1) the exposed wire must be large enough to fully cover 1 side of the sciatic nerve but not too large that would cause it to touch other soft tissues, 2) the 24 AWG is rather brittle and fails after several folding and unfolding cycles – so please check the integrity of the electrode or simply replace the electrode after each implantation, 3) the 24 AWG wire is stiff and has a tendency to either slide off the sciatic nerve or pull on the nerve, and 4) the cut ends should be fully out of the limb to prevent the exposed wire from contacting tissue and shorting the stimulator circuit.

**Fatigue effects over time.** During preliminary testing, we sought to minimize the time each animal was on the dynamometer. Earlier work suggested that 45 seconds was sufficient to minimize fatigue (Warren et al., 2004) to less than 3% over a series of 11 concentric contractions. Using our dynamometer with 15 second rest periods, we found that this shorter rest period resulted in similar fatigue across contractions (**Figure S2**). We found that the effects of fatigue were strongly linear (R2>0.999) with a relative decrease of 0.3% maximal plantar flexor torque each contraction. Because we randomized ankle angle during testing, we decided to not apply a fatigue correction. Further, based on the large effects we found (**Figure 2**), we do not expect this to have meaningful effects on our results. However, future work should explore fatigue correction when effect sizes are smaller.

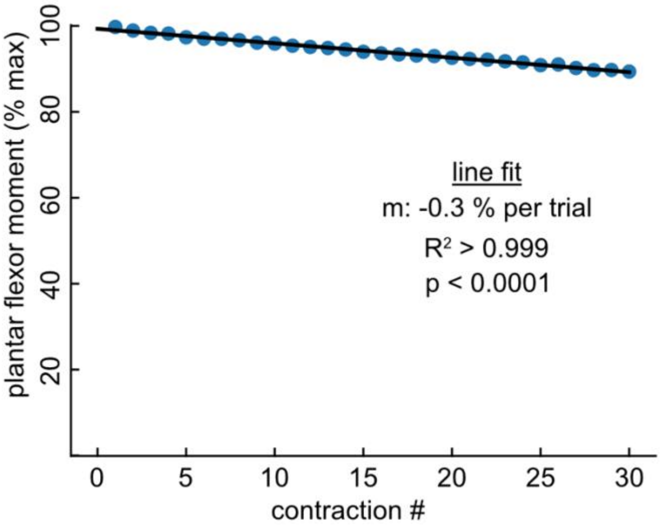


**Figure S2.** We performed 30 consecutive maximal contractions with the ankle held at 40 degrees dorsiflexion to test the effects of 15 second recovery on plantar flexor torque production. We found a strongly linear effect of repetitive contractions on plantar flexor torque. When normalizing to the peak torque (from contraction #1), we found that the relative fatigue was ~0.3% per contraction. Based on these findings, we determined that randomizing contraction order was sufficient. However, future work could implement a fatigue correction factor – especially if contractions at a constant ankle angle are repeated intermittently.

**Files available on GitHub.** All LabVIEW codes, STL files, and parts lists are freely available to download and use at https://github.com/joshrbaxter/rat_dynamometer.

**3D printed parts for securing animal to dynamometer.** We printed these parts on a Dremel 3D45 printer at 'high quality' settings. When printing with this specific printer, we used a 'raft' to ensure that the bottom of each part wasn't swollen that we have experienced when printing directly to the print bed. These parts should be able to be printed on other 3D printers with similar effectiveness. One point of note is the 'torque_cell_to_motor' part may benefit from a plastic table tie to secure the servo motor in the part.

**3D printed parts list.** Part files included in the supplemental: base_plate_torque_cell.stl, footplate_right.stl, knee_30degree_right.stl, nose_clip.stl, torque_cell_to_motor.stl. In your 3D printer layout software, mirror the foot_plate_right.stl and knee_30degree_right.stl to print left sided parts.

**LabVIEW files.** The control software runs on LabVIEW. Version used in this study was ratometer_beta0.5.vi. Additional functions required for dynamometer to function: analog_read.vi, build_digitimer_trigger.vi, build_protocol.vi, calculate_servo_PWM.vi, save_to_tdms.vi, trigger_digitimer.vi.

**Calibrating torque cell.** We calibrated the torque cell by hanging a known weight (9.32N) on 4 locations along the length of the servomotor arm. These paired screw holes were 12mm and 20mm from the servomotor spindle. We confirmed that the torque cell is linear ($R2 > 0.999$) and used the line fit parameters (m, b) in the LabVIEW program to transform torque cell voltages into torque measurements (N mm). Note, the torque cell conditioner has small set screws to adjust the span (measurement range) and zero (voltage value when torque is zero). During the calibration process, we set the span to cover approximately 650 N mm in both directions.

**Surgical Approach to the Sciatic Nerve Dissection SOP**

**Baxter Lab – see partslist.xlsx – Surgery parts sheet for tool list**

| | |
|---|---|
| 1) Shave the hindlimbs of the animal. Position the animal with both front and hind limbs extended. Secure limbs to table with tape. | 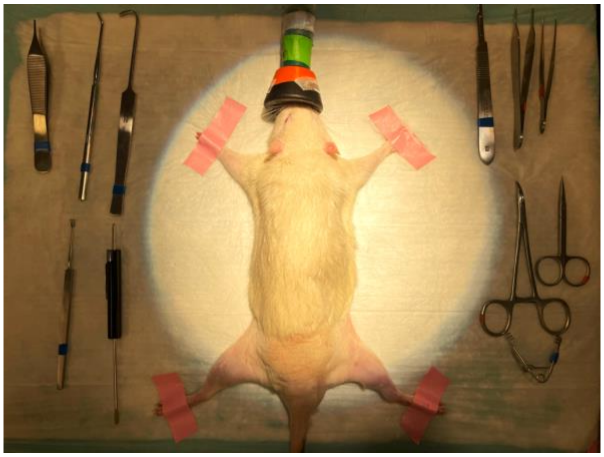 |
| 2) Identify location of femur with palpation of limb with forceps. Make an incision parallel with/just caudal to the femur at the level of the mid-femur; This can be done by gently lifting the skin and cutting with scissors Clear out fascia and undermine surrounding subcutaneous tissues as needed. | 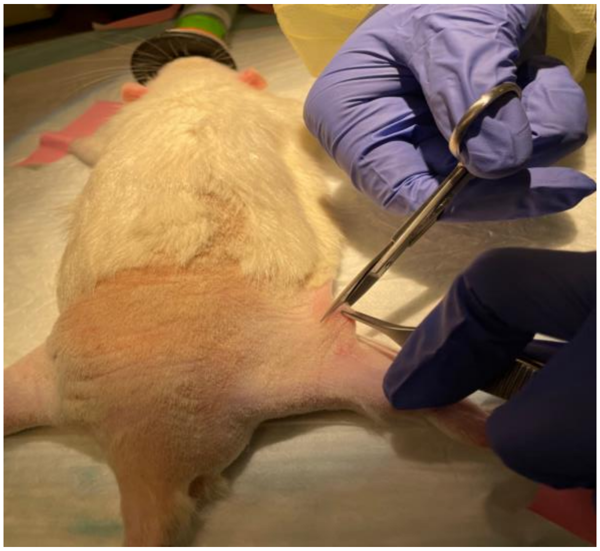 |
| 3) Identify white linear fascia separating adjacent muscle bellies | 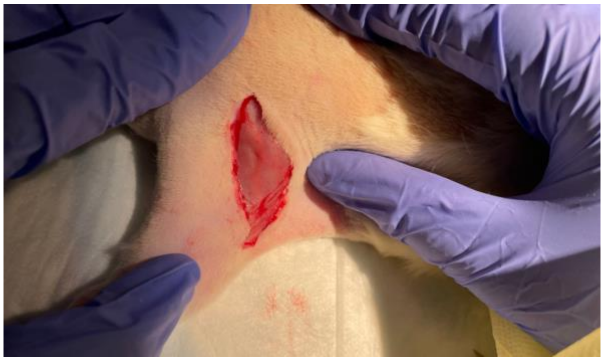 |

| | |
|---|---|
| 4) Using a scissor, make a small opening along the white linear fascia. Since the area beneath is hollow, this dissection should go clean and smooth with minimal bleeding. | 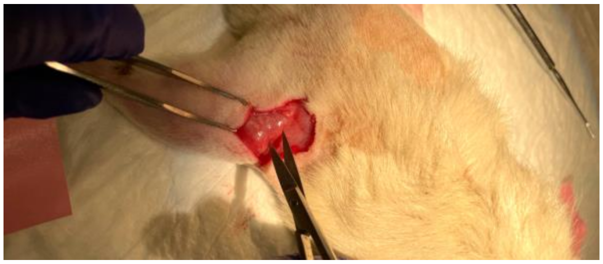 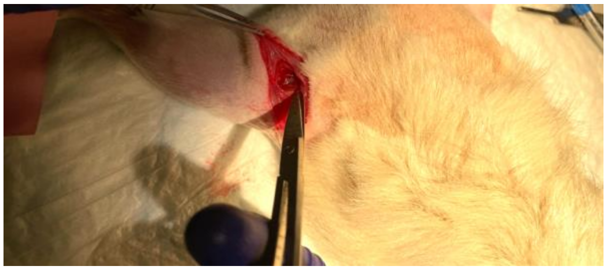 |
| 5) Once a small opening is made, expand this window using curved mosquito forceps. This will create a cavity large enough to clear the connective tissues around the sciatic nerve, providing space for electrode placement. | 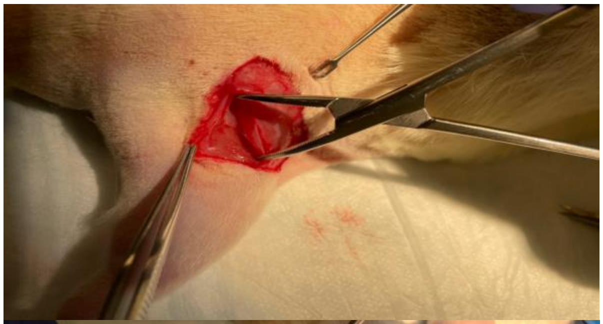 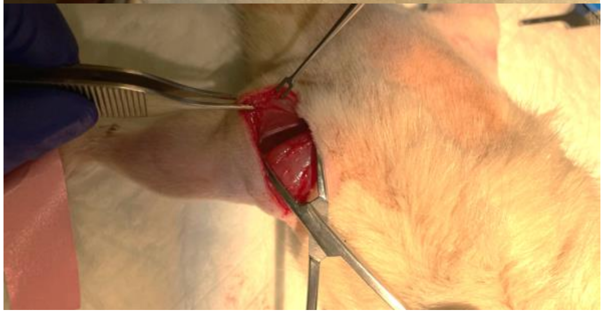 |

| | |
|---|---|
| 6) Identify the sciatic nerve sitting between these muscle bellies; Clear off fascia with blunt dissection with scissors as needed while ensuring that the surrounding nerve endings are not cut or damaged. | 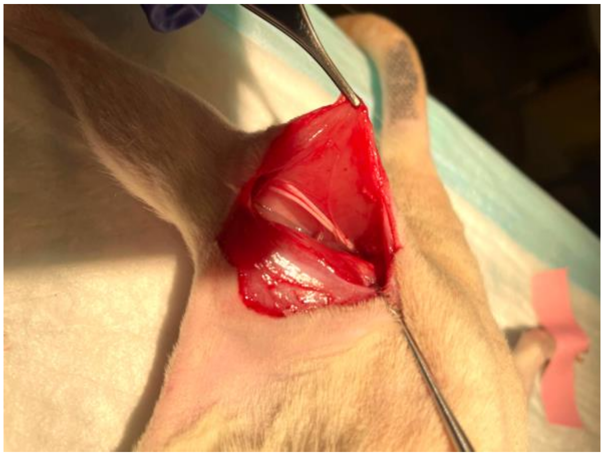 |
| 7) Glide a seeker with bent-end along the sciatic nerve to isolate it. | 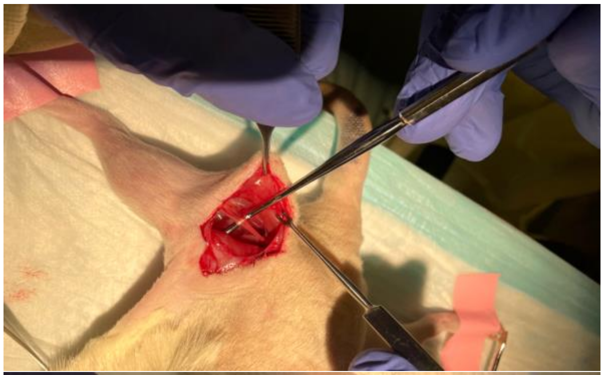 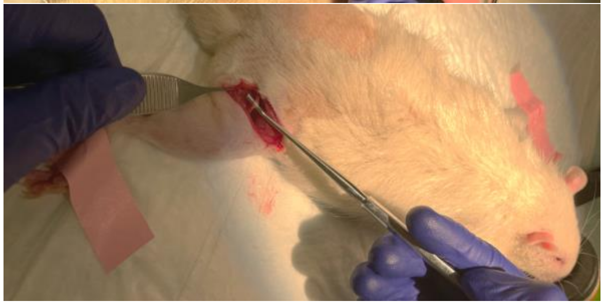 |
| 8) Once the sciatic nerve is fully isolated, eject .2-.3mL of mineral oil to keep it hydrated. Repeat the process a couple of times as needed throughout the dynamometer process. | 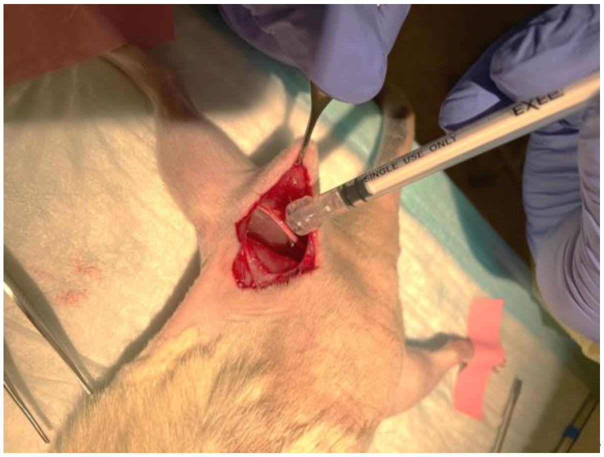  |

**Reminders:**

- Ensure that limbs are secured to the table with tape. This will help identify location of femur and prevent slippage during the blunt dissection.
- Avoid cutting the surrounding nerves whenever an incision is made to minimize bleeding
- Pour warmed mineral oil into the cavity to keep the nerve moist throughout the data collection

**Rat Dynamometer Software Guide**

**Baxter Lab - see partslist.xlsx, Dynamometer parts sheet for parts**

**Requisites:**

LabVIEW License

NI Data Acquisition Board - USB 6001 used in this system

Arduino - Rev 3 - 5V logic

Install Linx and upload to Arduino that controls servo motor

Hardware: Including DAQ, torque cell, amplifier, fixtures - see partslist.xlsx and STL folder

**Ratometer:**

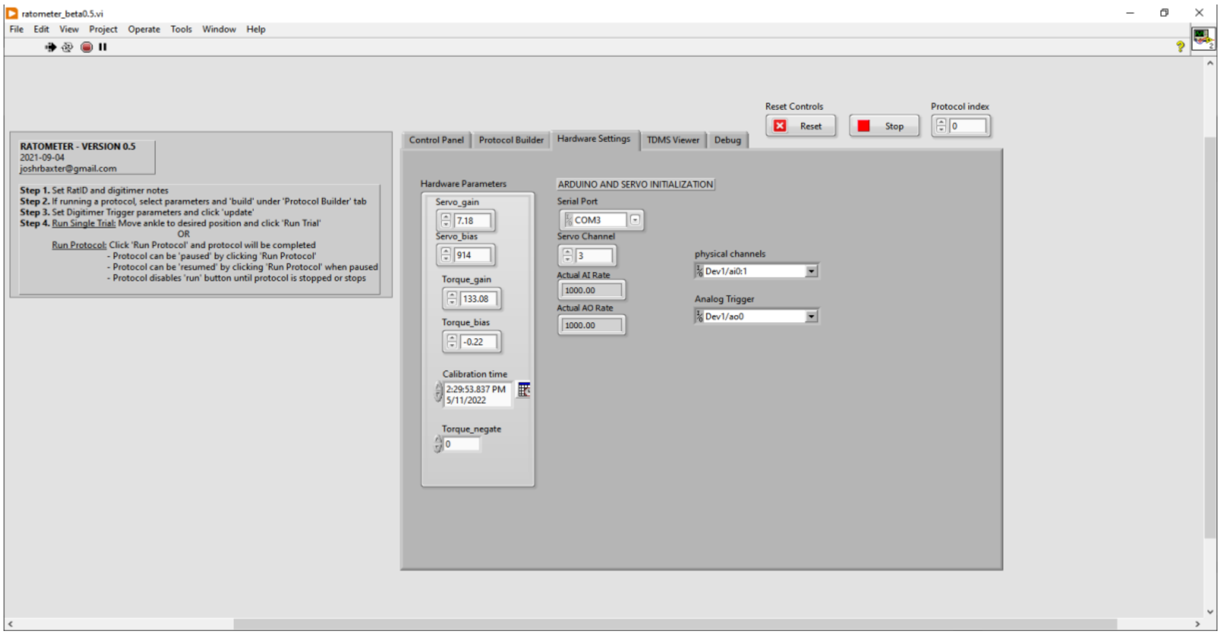


**Hardware Settings:** Before running the program, visit the "Hardware Settings" tab to set and define each parameter based on the user's set up.

**Reminder:**

- Before clicking the “Run” button located in the “Control Panel” tab, the user needs to visit the “Hardware Settings” tab and update the “Serial Port” as this parameter varies depending on the user’s setup.
- Torque gain and bias should be defined based on the type of torque cell that is being used.
- Servo gain and bias should also be defined based on the user's set up.
- Calibration time should be updated every time set up is calibrated.
- “Torque negate” should be left alone

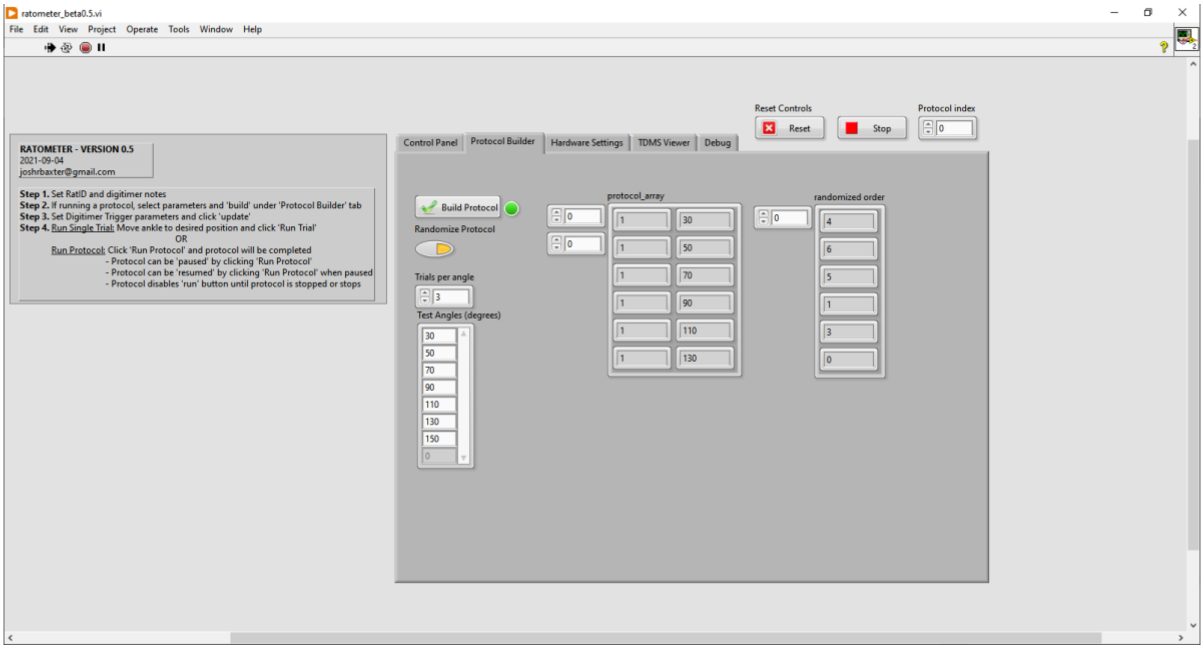


**Protocol Builder Tab:** Allows the user to set ankle positions, number of trials, and time of rest for each contraction set. It also allows users to randomize the order of each ankle position to eliminate testing sequence bias and to minimize the effects of fatigue.

**Reminder:**

- Before running the program, visit the “Protocol Builder” tab to build a protocol and randomize the order of each ankle position. Remember to click “reset” before each testing session begins.

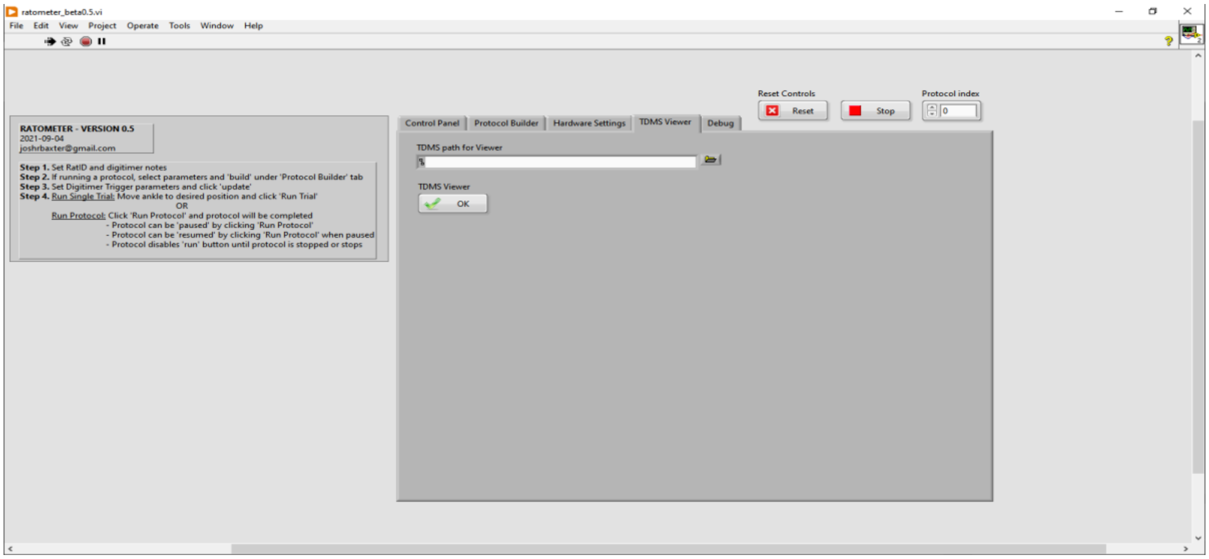


**TDMS Viewer Tab:** Used to view and locate existing experimental data.

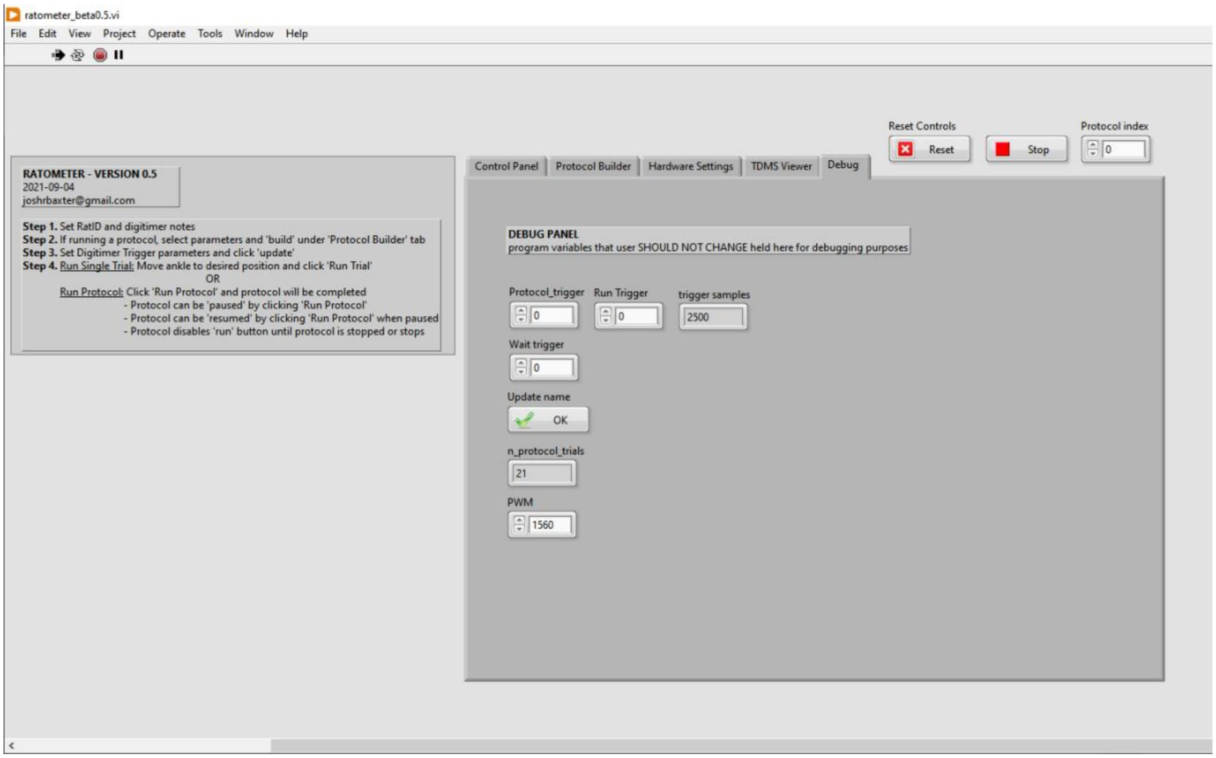


**Debug Tab:** A placeholder for variables that should not be changed by the user for debugging purposes.

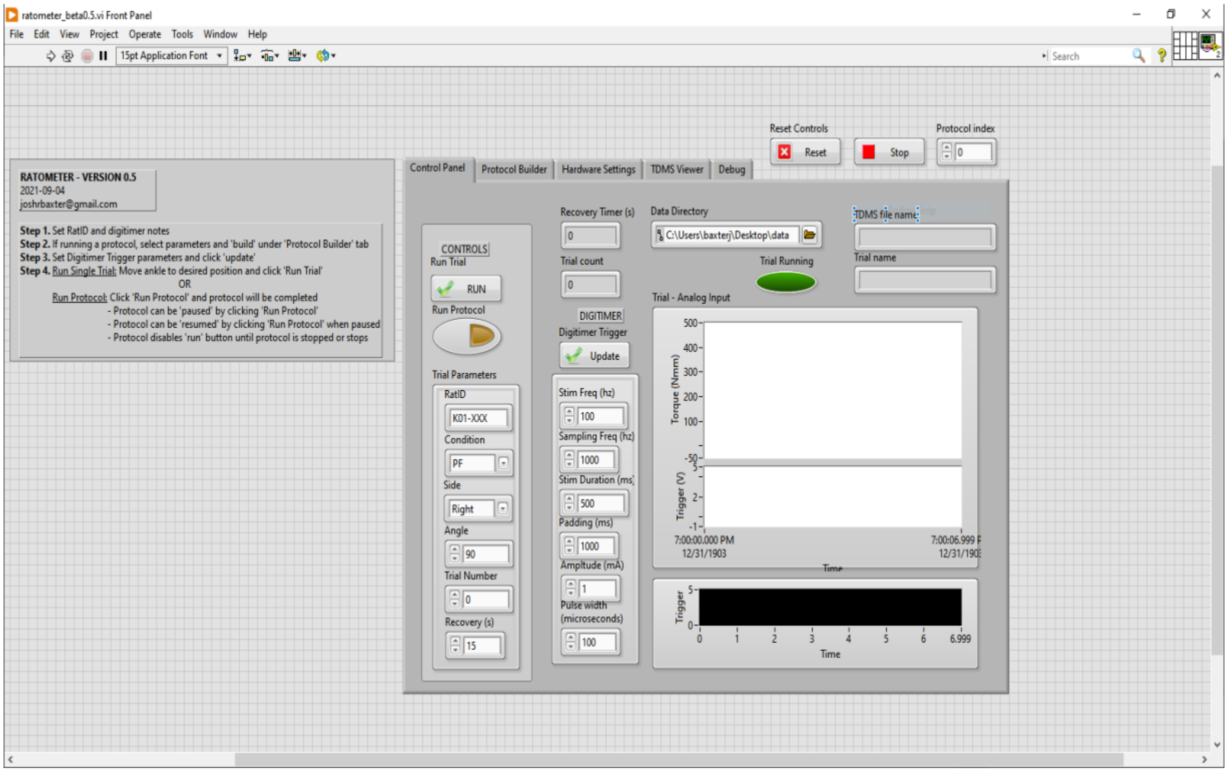


**Control Panel:** Location in which the user can run and save testing sessions.

**Reminder:**

- Before running the protocol, ensure that the data will be saved in the desired location under “Data Directory.” The Rat-ID and leg side should also be marked appropriately.
- Next, adjust the ankle angle under “Angle” to ensure that the foot-plate can freely rotate without bumping into other dynamometer components such as the load-cell.
- Under “Condition,” run a “Max” test at 90 degrees. The torque result should be around 250-300 Nmm. If the user obtained a lower torque measurement, double-check the position of the nerve cuffs to ensure correct placement. The cuffs should be firmly holding the sciatic nerve with the surrounding tissues cleared of sight. Adjust the “Amplitude” from 1 to 2 mA or vice versa as needed.
- Change the condition to “PF” and stimulation duration to 500 ms before running. Refer to the picture above for the standardized set-up. Remember to always click “Update” whenever a change is made. Once the parameters are set, the user may begin running the program.
- After the session is finished, obtain a fatigue data by setting the condition

| |
|---|
| to “Fatigue” and stimulation duration to “5000 ms” |

**Block Diagram:**

*To be used as a reference

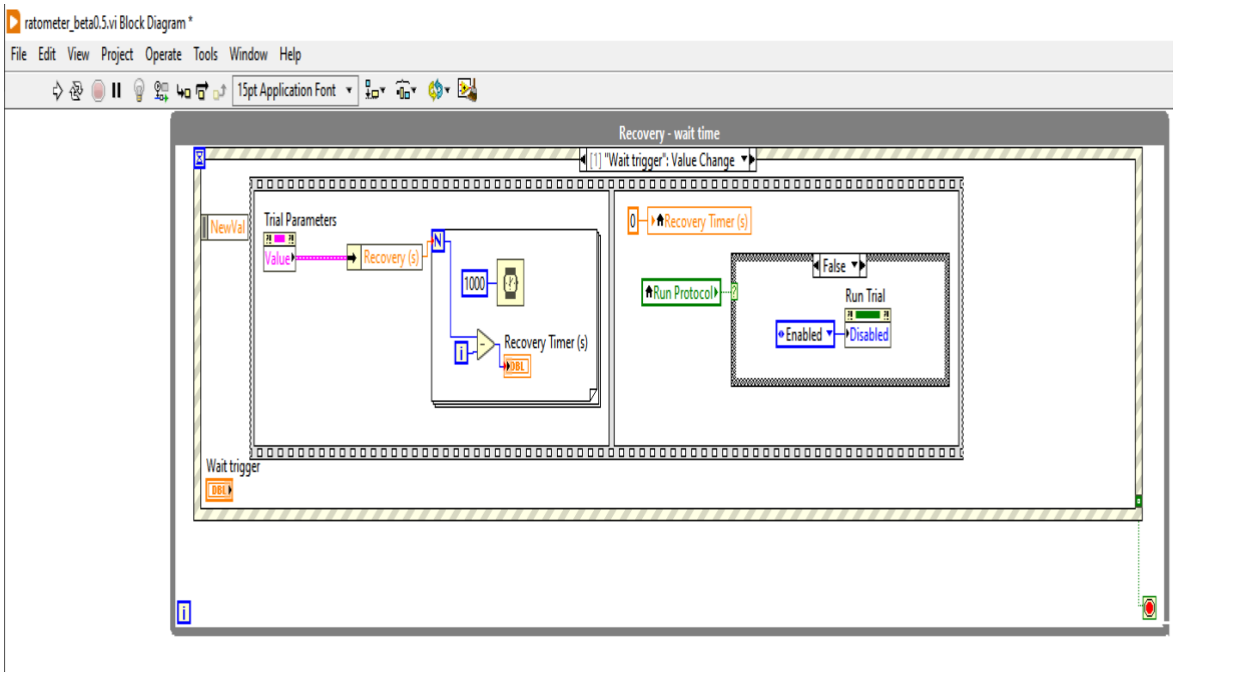

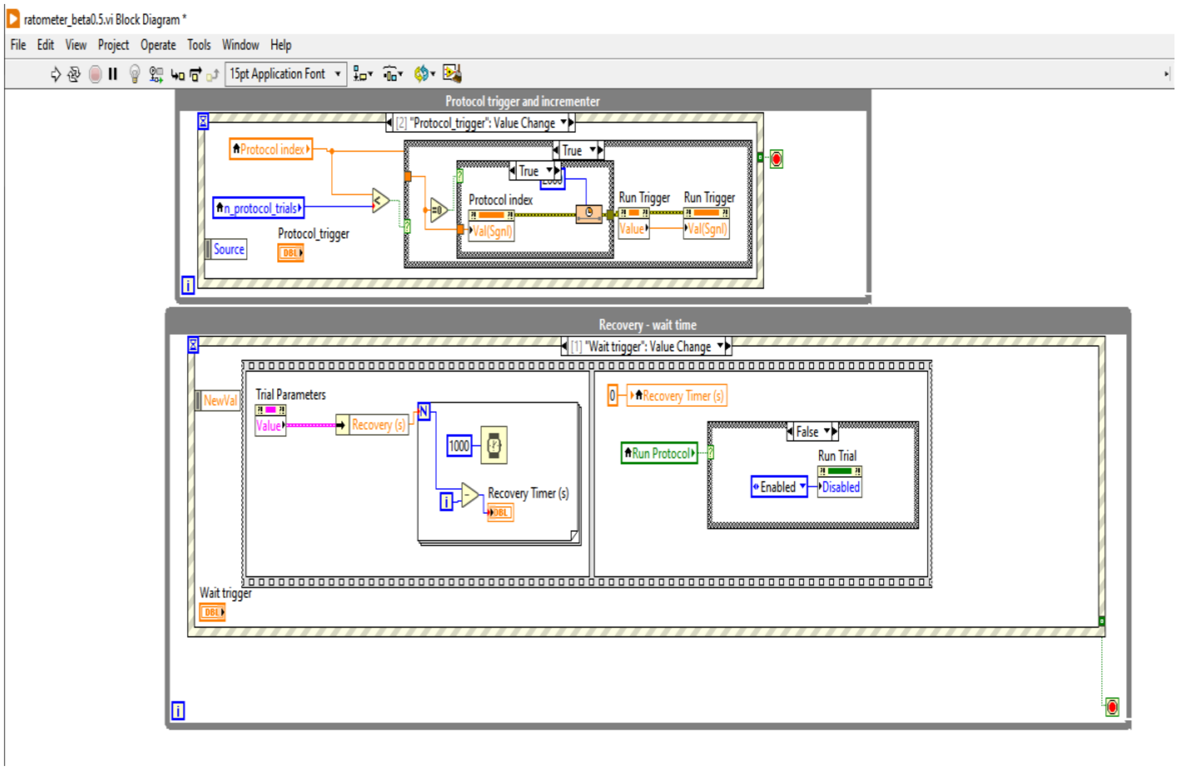

ratometer_beta0.5.vi Block Diagram *
File Edit View Project Operate Tools Window Help
15pt Application Font
Protocol trigger and incrementer
[2] "Protocol_trigger": Value Change
Protocol index
n_protocol_trials
Source
Protocol_trigger
True
Protocol index
Val(Sgnl)
Run Trigger
Value
Run Trigger
Val(Sgnl)
Recovery - wait time
[1] "Wait trigger": Value Change
NewVal
Trial Parameters
Value
Recovery (s)
1000
Recovery Timer (s)
0
Recovery Timer (s)
Run Protocol
False
Run Trial
Enabled
Disabled
Wait trigger


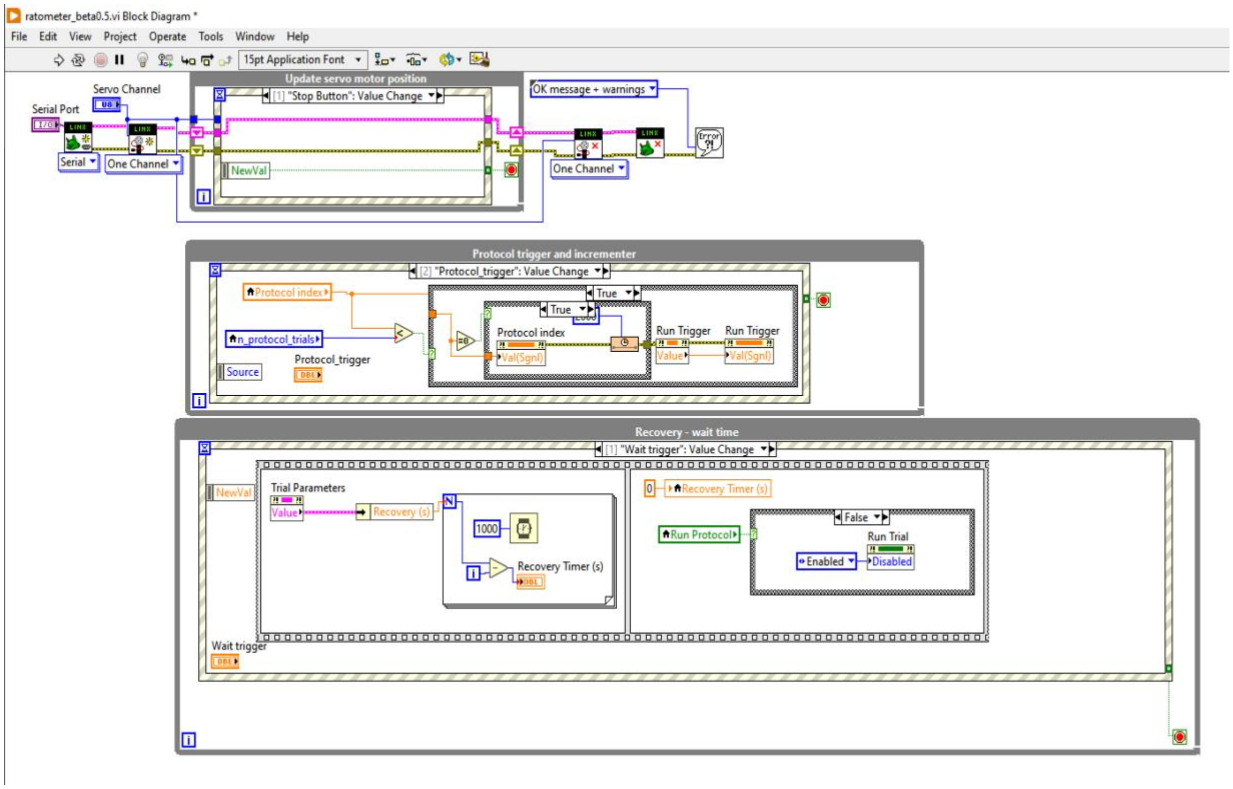

ratometer_beta0.5.vi Block Diagram *
File Edit View Project Operate Tools Window Help
15pt Application Font
Servo Channel
Serial Port
Serial
One Channel
Update servo motor position
[1] "Stop Button": Value Change
NewVal
OK message + warnings
One Channel
Protocol trigger and incrementer
[2] "Protocol_trigger": Value Change
Protocol index
n_protocol_trials
Source
Protocol_trigger
True
Protocol index
Val(Sgnl)
Run Trigger
Value
Run Trigger
Val(Sgnl)
Recovery - wait time
[1] "Wait trigger": Value Change
NewVal
Trial Parameters
Value
Recovery (s)
1000
Recovery Timer (s)
0
Recovery Timer (s)
Run Protocol
False
Run Trial
Enabled
Disabled
Wait trigger

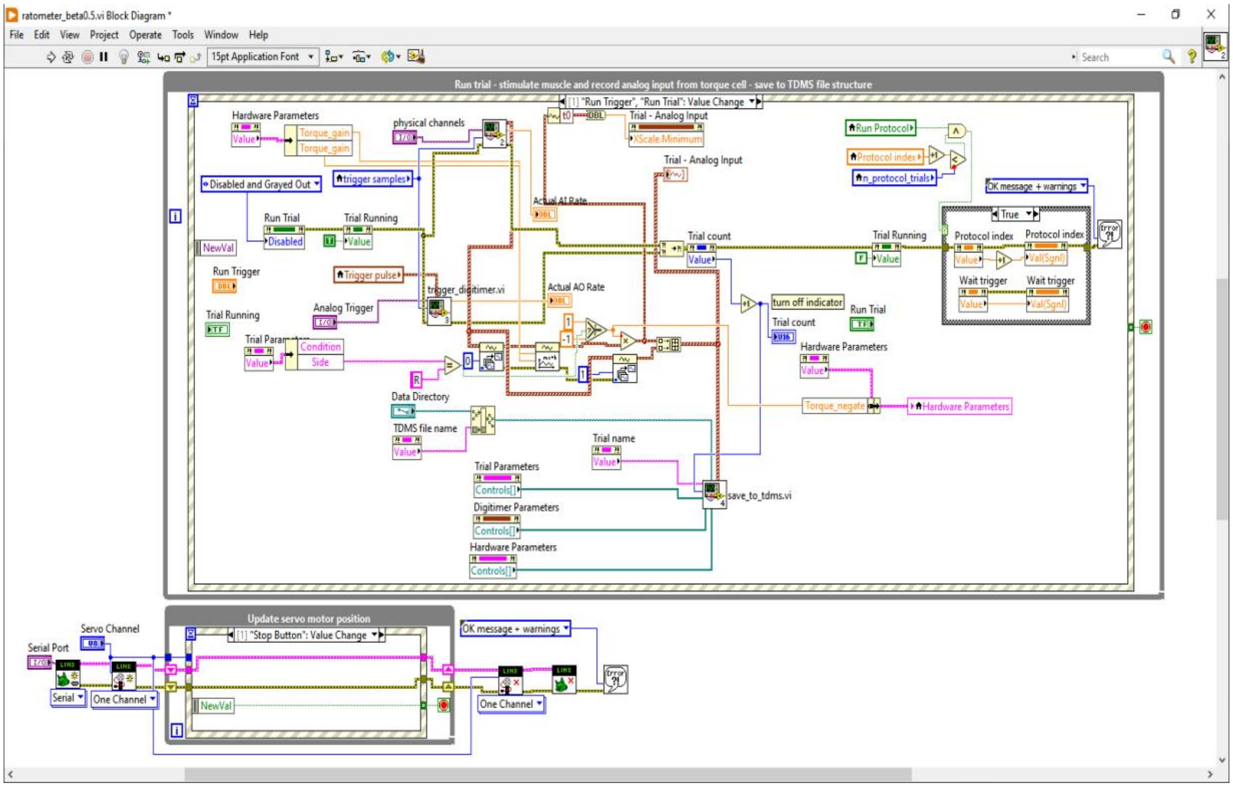
ratometer_beta0.5.vi Block Diagram *
File Edit View Project Operate Tools Window Help
15pt Application Font
Search
Run trial - stimulate muscle and record analog input from torque cell - save to TDMS file structure
[1] "Run Trigger", "Run Trial": Value Change
Hardware Parameters
Torque_gain
physical channels
Trial - Analog Input
XScale.Minimum
Run Protocol
Protocol index
n_protocol_trials
OK message + warnings
Disabled and Grayed Out
trigger samples
Actual AI Rate
Run Trial
Trial Running
Disabled
NewVal
Run Trigger
Trigger pulse
trigger_digitimer.vi
Actual AO Rate
Trial count
True
Wait trigger
turn off indicator
Analog Trigger
Condition
Side
Data Directory
TDMS file name
Torque_negate
Trial name
Trial Parameters
Controls[]
Digitimer Parameters
save_to_tdms.vi
Update servo motor position
Servo Channel
Serial Port
[1] "Stop Button": Value Change
Serial
One Channel

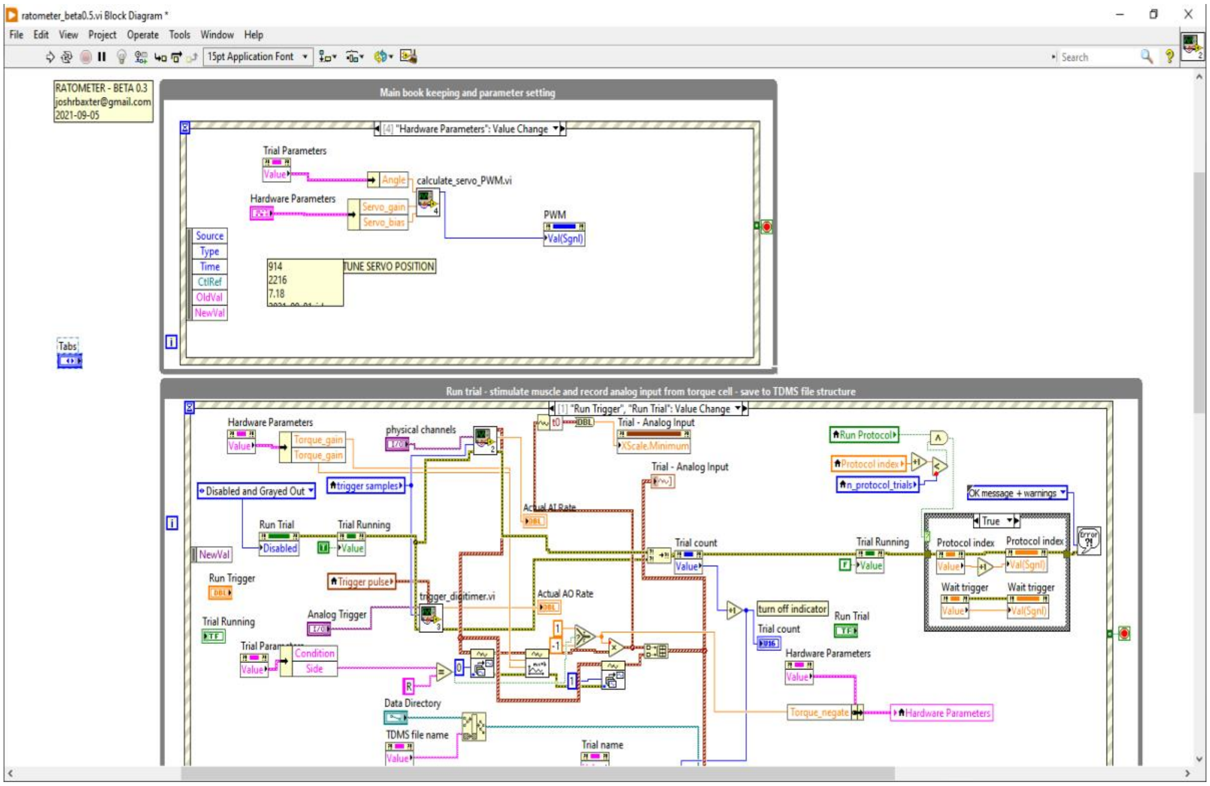
ratometer_beta0.5.vi Block Diagram *
File Edit View Project Operate Tools Window Help
15pt Application Font
Search
RATOMETER - BETA 0.3

2021-09-05
Main book keeping and parameter setting
[4] "Hardware Parameters": Value Change
Trial Parameters
Angle
calculate_servo_PWM.vi
Hardware Parameters
Servo_gain
Servo_bias
PWM
Val(Sgnl)
Source
Type
Time
CtlRef
OldVal
NewVal
914
2216
7.18
TUNE SERVO POSITION
Tabs
Run trial - stimulate muscle and record analog input from torque cell - save to TDMS file structure
[1] "Run Trigger", "Run Trial": Value Change
physical channels
Trial - Analog Input
Run Protocol
Protocol index
n_protocol_trials
OK message + warnings
Disabled and Grayed Out
trigger samples
Actual AI Rate
Run Trial
Trial Running
Trial count
Wait trigger
Run Trigger
Trigger pulse
trigger_digitimer.vi
Actual AO Rate
turn off indicator
Analog Trigger
Condition
Side
Data Directory
TDMS file name
Trial name
Torque_negate
